# The Evolutionary Flexibility of the Drosophila Circadian Clock: Network Constraints or Adaptive Freedom?

**DOI:** 10.1101/2025.04.02.646778

**Authors:** Leo Douglas Creasey, Eran Tauber

## Abstract

The study of network evolution is critical to understanding how complex biological processes arise and adapt over time. Protein networks, composed of interacting components, can exhibit varying degrees of conservation and flexibility, enabling organisms to fine-tune their responses to environmental changes. Using the circadian clock system in Drosophila as a case study, we explore how such networks evolve. We leverage the recently published 101 Drosophilidae genome project to analyze the evolution and co-evolution of 11 core clock proteins across 65 species spanning about 60 million years of evolution. A sliding window analysis of coding regions reveals substantial heterogeneity in nucleotide divergence, with *Clk* and *per* exhibiting high divergence, whereas *Pdp1* and *sgg* show virtually no evolutionary change. Additionally, we assessed interdependent amino acid evolution across different proteins, identifying 67 co-evolving site pairs, primarily between CLK-PER, CLK-CWO, and SGG-PER. Using codon-based models of evolution we found four genes (*cwo, jet, per*, and *sgg*) showing evidence of positive selection. Since several clock proteins are pleiotropic, we tested whether their multifunctionality influences their evolutionary constraints. Using alternative approaches to assess pleiotropy, we found no significant correlation between pleiotropy and the non-synonymous substitution rate (Ka) in 440 *Drosophila* proteins, including circadian clock ones. Overall, our findings suggest that the circadian clock network does not impose strong constraints on the evolution of its components. This flexibility may facilitate species-specific adaptation of the clock and allow the pleiotropic functions of clock proteins.

## 1 Introduction

In multicellular organisms such as animals and plants, the number of phenotypic states far exceeds the number of proteins within their proteomes. This expansion of phenotypic diversity has been achieved through the evolution of complex gene and protein networks [1]. Mechanisms such as gene duplications enable proteins to acquire novel functions, thereby increasing the diversity of interactions within biological systems[2]. Gene duplications not only expand the protein network but also promote the development of new phenotypes, as duplicated genes diverge in function over time. Furthermore, mutations in existing proteins—referred to as link dynamics—serve as a dominant evolutionary force shaping network structure [3]. The interplay between gene duplications, link dynamics, and stochastic variations in gene expression contributes to the rich diversity of protein networks. These processes illustrate how gene networks evolve to generate new phenotypes and enhance biological diversity through structural and functional innovations.

The circadian clock (Latin for circa “around” and diem “day”) is an endogenous pacemaker that drives diel physiological and biochemical rhythms. The circadian system allows organisms to track the predictable daily changes in their environment, such as light and temperature, resulting from Earth’s self-rotation. This ability to anticipate environmental changes and synchronize behavior with optimal times of day provides a strong adaptive advantage. For example, the movement of sunflowers toward the east to face the rising sun and honeybee visits to specific flowers at particular times are both governed by circadian rhythms [4].

Much of the understanding of the molecular circuit underpinning the circadian system has emerged from research on the fruit fly *Drosophila melanogaster*. The first clock gene to be discovered in the fruit fly was *period* (*per*) which as the following genes that have been discovered, is highly conserved in all other animal groups, including humans.

The *Drosophila* circadian clock is composed of two interlocking negativefeedback loops (Figure 1)[5]. The core loop begins with the dimerization of CLOCK (CLK) and CYCLE (CYC), which bind to the E-box domains of the *period* (per) and *timeless* (tim) genes, promoting their transcription. The resulting mRNA is processed and translated, and the PER and TIM proteins dimerize and enter the nucleus. Once inside the nucleus, the PER-TIM complex inhibits CLK-CYC and suppresses its own transcription. This leads to a decrease in *per* and *tim* mRNA levels, followed by a reduction in the corresponding protein levels, which terminates the inhibition of CLK-CYC, thus allowing the initiation of a new diurnal cycle. The PER-TIM autorepression is further reinforced by CLOCK-WORK ORANGE (CWO), which binds to E-boxes to maintain the PER-TIM complex on the CLK-CYC-bound DNA [6].

**Fig. 1.**
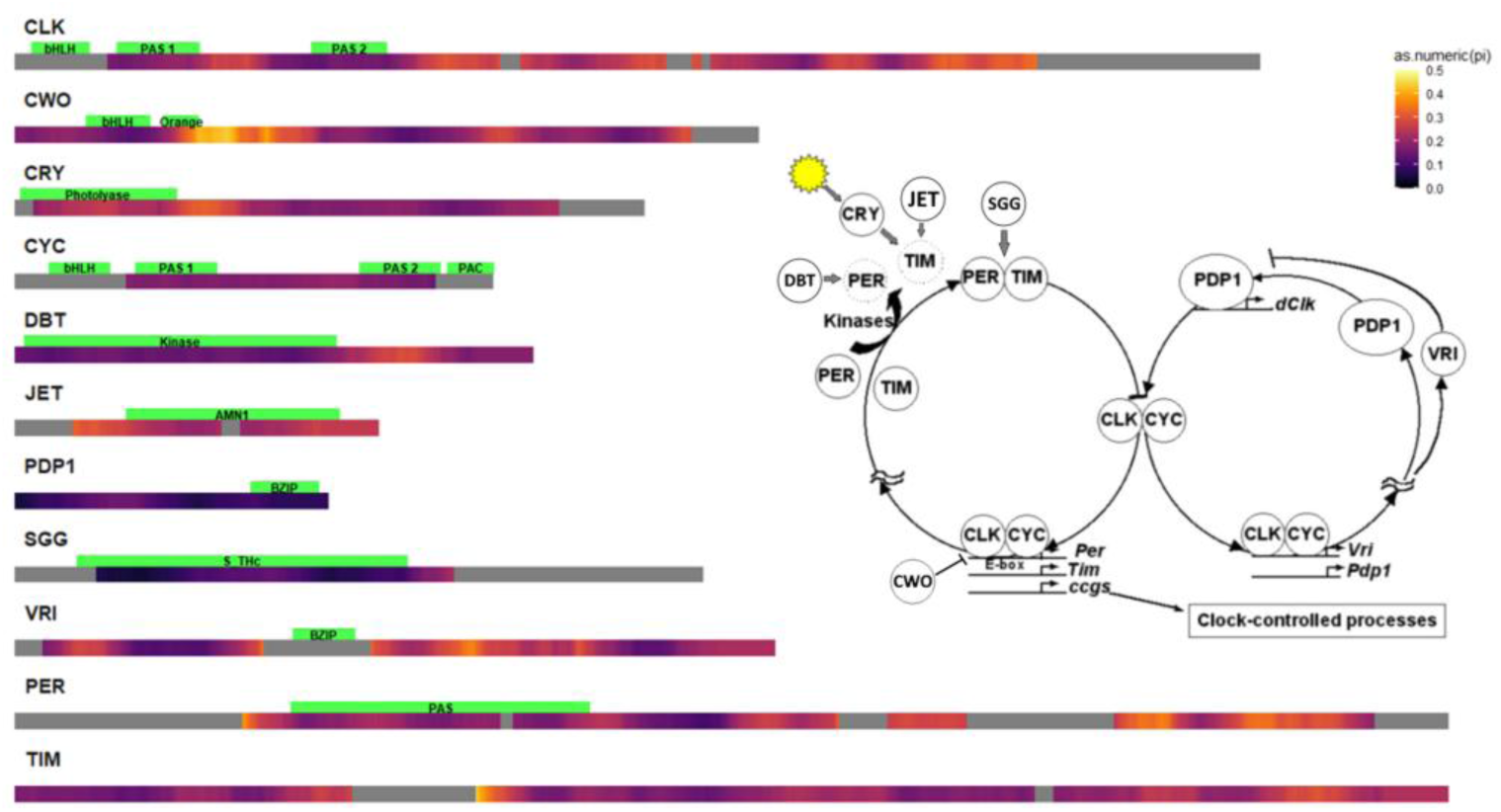
Nucleotide divergence of circadian clock genes across Drosophila species. Heatmap showing relative pairwise genetic divergence (Dxy) using a 150 bp sliding window. Inset: Schematic representation of the circadian clock molecular circuit, illustrating the core feedback loops and interactions among clock genes.

The timing of the circadian cycle is regulated by the phosphorylation of PER and TIM proteins, a process carried out by kinases such as DOUBLETIME (DBT) and SHAGGY (SGG), which influence their stability and nuclear localization. In addition to the PER-TIM feedback loop, a secondary loop operates in parallel. In this loop, Vrille (VRI) suppresses *Clk* gene expression, while PAR-domain protein 1ε (PDP1ε) enhances it. The output of the oscillator is mediated by proteins such as the pigment-dispersing factor (PDF) neuropeptide and its corresponding receptor (PDFR), which play a key role in synchronizing the circadian clock with the organism’s behavioral rhythms.

The entrainment of the circadian period to the external 24-hour light-dark cycle is mediated by light-dependent degradation of TIM, an interaction involving CRYPTOCHROME (CRY), a blue light photoreceptor [7], and JETLAG (JET), which further modulates the synchronization of the clock to the environmental light-dark cycle.

The circadian clock network is particularly interesting because several clock proteins are pleiotropic, serving roles beyond circadian regulation. For instance, TIM is involved in immune responses [8], CRY and PDF influence geotaxis [9], and SGG participates in the Wnt signaling pathway, essential for development and organogenesis [10]. Given their broad functions, pleiotropic genes are subject to purifying selection, which reduces genetic variation [11]. However, pleiotropy can also lead to balancing selection, maintaining genetic diversity by favoring different alleles for their effects on distinct traits [12].

Previous studies have explored genetic variation in clock genes across different *Drosophila* species, demonstrating how the evolution of these genes drives molecular adaptations specific to the species’ ecological niches [13]. These studies have typically focused on the evolution of individual genes within the circadian clock, often in isolation. However, these genes operate as part of a highly interconnected network, and the evolution of the clock system has not been studied as a whole network of interacting proteins. The recent genome sequencing of 93 *Drosophila* species [14] allow for a comprehensive analysis of the evolution of circadian clock genes across multiple species. These species represent approximately 60 million years of evolutionary history and span 14 subgroups within *Drosophila* and *Sophophora* [15]. Here, we analyzed this dataset and identified the orthologous clock proteins across all these species, providing a unique opportunity to examine the evolution of circadian clock genes not in isolation, but as part of a broader, interconnected gene network. One of the key questions addressed in this study is how do pleiotropic genes within the circadian clock network evolve, and how do their interactions with other genes in the network contribute to the broader evolutionary dynamics of gene networks.

## 2 Methods

### 2.1 Gene Prediction and protein alignment

The 101 Drosophila genome project consisted of 101 genome scaffolds [14], lacking gene annotation. To identify the Drosophila 11 clock proteins (Fig. 1) we used AUGUSTUS software for gene prediction [16]. The search for the orthologous gene was facilitated using *D. melanogaster* sequences. These sequences were used in a BLAST search against a database of every genome from the 101 Drosophila genomes project. These BLAST fragments were then used to search for the predicted genes from the respective species. The collected sequences are aligned using MAFFTv7.490 [17]. These alignments are then scored using Alistat [18] where species sequences that had less than 40% consensus were removed. The sequences were converted to amino acid sequences, and were re-aligned using the codon information. Finally, the alignments were adjusted by visual inspection. The final alignment included sequences from 65 different Drosophila species. The predicted gene files, and the pipeline scripts are available at Zenodo (DOI available upon publication), and GitHub (https://tinyurl.com/25xmsbkt) respectively.

### 2.2 DNA variation analysis

To assess interspecific genetic variation, we calculated the average pairwise nucleotide divergence across Drosophila species, analogous to the π statistic. To that end, we used the *pegas* R package [19]. To minimize the sites with gaps in the data, we carried out pruning of the alignments, where sequences with a high percentage of gaps (over 5% of the total sequence length, or over 100 gap bases total) were removed. Using a custom code, the polymorphism was computed for a sliding window of a 180 bp window length, and 6 bp step size. The output of the analysis was exported as text files and heatmap plots depicting polymorphism along the coding regions were created.

We carried out a non-synonymous-to-synonymous substitution ratio test (the *K*_A_/*K*_S_ test, ω) using a custom-made script and the SeqinR R package [20]. The script runs a sliding window of 150 bp and a 6 bp step. This analysis provides a visual representation of the variation in selection pressure, with ω < 1 indicating purifying selection, ω ≈ 1 indicating neutral evolution, and ω > 1 suggesting positive selection. Graphs were made using the R software.

To test for the signature of negative purifying selection or accelerated evolution driven by positive Darwinian selection, we used the CodeML program from the PAML package [21]. We generated the models M0 (One-ratio model), M1 (Nearly neutral model), M2 (Positive selection model), M7 (Beta model), and M8 (Beta & ω > 1 model). We compared the models using a likelihood ratio test. A custom script was made to conduct the model comparison and parse the significant sites from the results files.

We used the CAPS 2.0 software [22] to predict coevolution both within genes (intra) and between genes (inter) at a significance level α = 0.05. A custom script pulled the relevant sites. Sites were rejected if they had a correlation of less than 0.5, or a bootstrap score of less than 0.95. The R package *circlize* was used to create the circular graphs [23].

To assess pleiotropy of *Drosophila* genes we used two distinct methods. The first utilized the DRoID interactome database [24]. A custom-written code screened the database files and counted the number of times a gene of interest interacted with other genes. The database files that were used included yeast-2-hybrid files and Flybase genetic interaction database (duplicate interactions were counted once).

The second method of quantifying pleiotropy was based on a gene of interest’s number of Gene Ontology (GO) terms. Only the highest-level terms of the GO hierarchy (’Biological Process’) were counted. The consistency of these methods was compared using R. The rate of non-silent substitutions (Ka) across species was calculated as described above, and the effect of pleiotropy was tested using (i) the gene-wide overall Ka (ii) the maximum Ka (Ka^*^) from the sliding window analysis. A linear model was carried out in R using the MASS library with a Box-Cox transformation to stabilize the variance of the Ka data.

### 2.3 Phylogenetic trees and Tanglegrams

The DNA alignments were used for generating a distance matrix using the “K80” model using the *ape* R library [25]. A Neighbour-Joining tree was created using the default arguments and converted to an ultra-metric tree using the *chronos()* function. The ‘mirror-trees’ (tanglegrams) of pairs of clock proteins were generated using the *dendextend* library [26], and their co-evolution was visually analaysed.

## 3 Results

### 3.1 Sequence divergence of circadian clock genes

The sliding window analysis of the average pairwise nucleotide divergence (Dxy) of circadian clock genes revealed substantial variation between and within genes (Figure 1). Among the different genes, *cwo* and *tim* exhibited the highest levels of variability, with *tim* displaying the widest range of Dxy values, followed closely by *cwo*. These genes showed substantial fluctuations across their coding regions, suggesting a high degree of localized molecular polymorphism. *Clk* and *per* also exhibited relatively high levels of diversity, with mean Dxy values around 0.23 and broad distributions spanning 0.105– 0.379 and 0.137–0.344, respectively.

In contrast, *Pdp1* and *sgg* were the most conserved genes in terms of nucleotide divergence. *Pdp1* had the lowest mean Dxy (0.10) with a narrow distribution ranging from 0.051 to 0.155. Similarly, *sgg* exhibited a low mean Dxy (0.10) and the smallest minimum value (0.04), indicative of strong sequence conservation. *cyc* and *dbt* also demonstrated lower diversity, with mean divergence values around 0.17 and relatively stable distributions.

Variation within genes showed different patterns, with some genes such as *jet* and *cry* displaying relatively uniform levels of polymorphism across their length, whereas others, particularly *cwo* and *tim*, showed distinct regions of elevated diversity interspersed with conserved stretches. *Clk, per*, and *vri* exhibited intermediate levels of variation, with notable peaks and troughs in diversity along their coding sequences.

### 3.2 Inter-dependent evolution of amino acids in clock proteins

The analysis of coevolution between clock proteins using CAPS identified 67 significant coevolving sites (Figure 2). A total of 20 co-evolving site pairs were detected between CLK and PER, 11 site pairs between CLK and CWO, 14 sites between SGG and PER, 8 site pairs between PER and CWO, and 5 pairs between PER and CYC. A single pair of coevolving sites was identified between CRY and CWO, TIM and SGG, PER and JET, CWO and PDP1, CWO and CRY. Among these interactions, some amino acid sites were involved in multiple co-evolving pairs, indicating potential shared functional constraints. A single site in CWO appeared in all the pair sites in CLK, PDP1 and PER. A single site in SGG was involved in interaction with PER, and CLK. The distribution of co-evolving sites was not uniform across the sequences, with some regions exhibiting a higher density of co-evolutionary links. Notably, no co-evolving sites were identified in protein pairs that are known to form heterodimers such as PER and TIM [27] or CLK and CYC [28]. The lack of apparent co-evolution between proteins (or domains within) that physically interact with each other is consistent with a recent study demonstrating that compensatory coevolution due to physical interaction contributes to evolutionary rate covariation is weak compared to the contribution of shared changes in selective pressure [29].

**Fig. 2.**
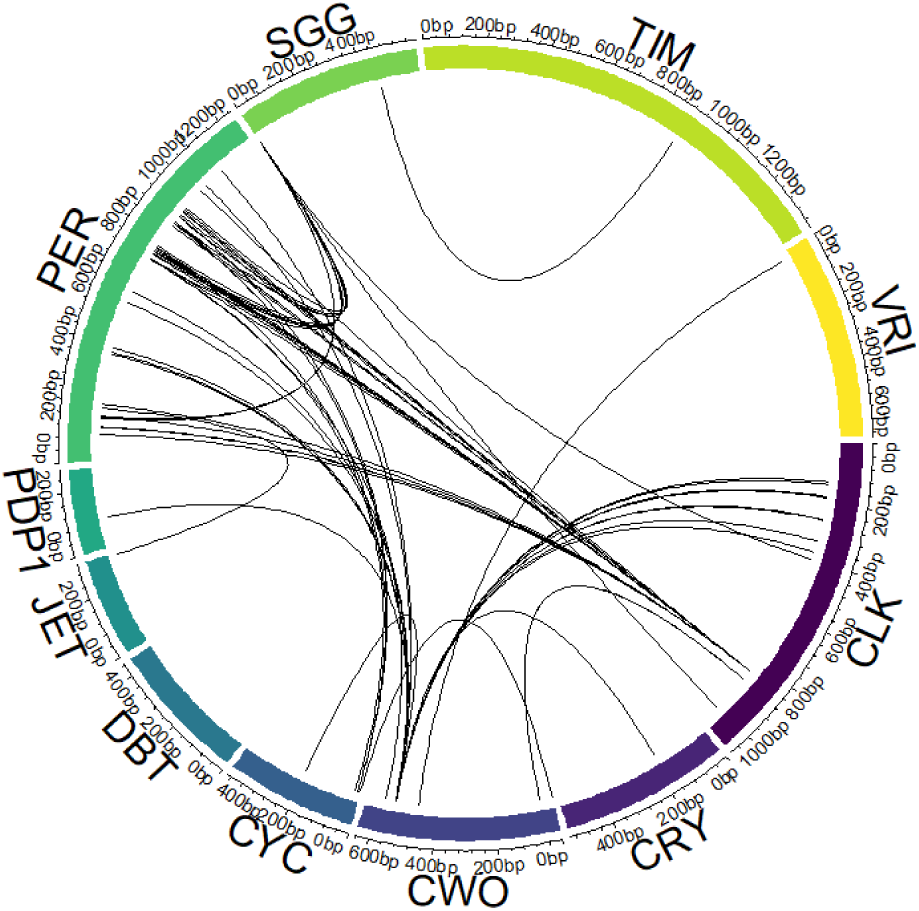
Co-evolving sites among circadian clock proteins. The circular plot shows significant co-evolving pairs of amino acids (p < 0.05).

As expected, the number of intra-protein coevolving sites is substantially higher than the number of inter-protein sites (Supplementary Figure 1). PER and TIM have the highest number of intra-protein site pairs (506 and 444, respectively), and PDP1 and DBT have the smallest number of intra-protein interactions (6 and 30).

### 3.5 Selection Analysis

To investigate patterns of selection pressure across the 11 circadian clock genes, we conducted a sliding window analysis to estimate the Ka/Ks ratio (ω) along the length of each gene (Figure 3). The majority of genes exhibited consistently low Ka/Ks ratios (ω << 1) throughout their lengths, indicative of strong purifying selection. These genes included *Clk, cry, cyc, dbt, pdp1, tim*, and *vri*. This suggests strong evolutionary constraint, indicating high conservation and a small number of changes. While no genes displayed Ka/Ks ratios exceeding unity, suggesting pervasive positive selection, some genes exhibited regions with slightly elevated Ka/Ks ratios compared to the baseline. For *per*, the sliding window analysis showed peaks in ω within certain regions along the length of the sequence. A trend toward an increasing ω was observed toward the 3’ end in *sgg*, potentially suggesting relaxed evolutionary selection in this part of the gene. Subtle peaks in Ka/Ks ratios were also apparent along the lengths of *cwo* and *jet*, hinting at localized selective pressures.

**Fig. 3.**
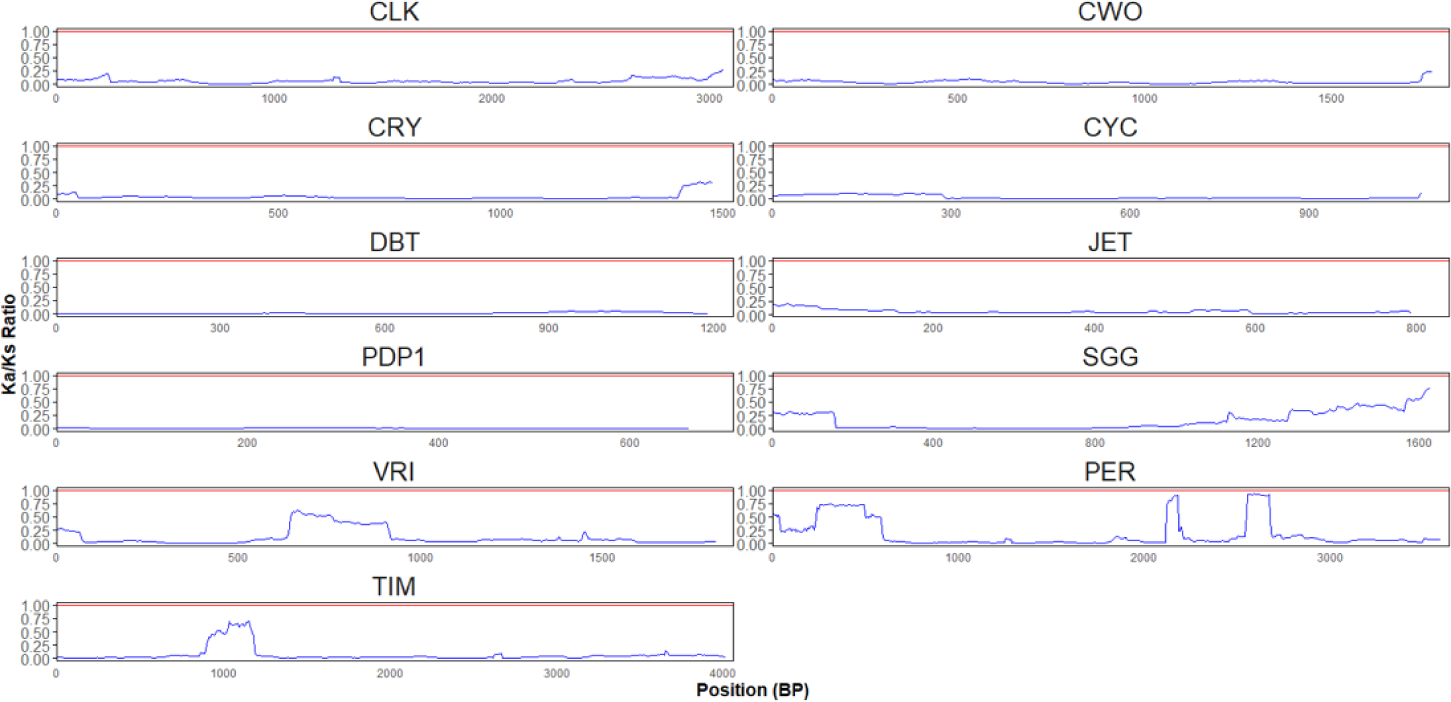
Comparative analysis of selection pressures across Drosophila circadian clock genes using sliding window analysis of Ka/Ks ratios. The ratio of non-synonymous to synonymous substitution rates (ω) was calculated to identify signatures of selection.

To rigorously test for evidence of site-specific selection, we employed codon-based maximum likelihood models implemented in the PAML package. Specifically, we compared models allowing for positive selection (M8) to those that do not (M7) using likelihood ratio tests (LRTs). The results of these analyses are summarized in Table 1.

**Table 1.** Likelihood ratio tests (LRTs) for positive selection in *Drosophila* circadian clock genes. The table summarizes the results of PAML analyses comparing different codon substitution models to detect positive selection. Each cell is color-coded to indicate the significance of the likelihood ratio test for the specified model comparison. Blue cells indicate a statistically significant result (p < 0.05), supporting positive selection, while magenta cells indicate a non-significant result. The model comparisons shown are M0vsM1 (One Ratio vs. Multiple Ratio), M1vs M2 (Nearly Neutral vs. Positive Selection), and M7 vs M8 (Beta vs. Beta & Positive).

|  | M0 vs M1 | M1 vs M2 | M7 vs M8 |
| --- | --- | --- | --- |
| CLK | Blue | Magenta | Magenta |
| CRY | Blue | Magenta | Magenta |
| CWO | Blue | Magenta | Blue |
| CYC | Blue | Magenta | Magenta |
| DBT | Blue | Magenta | Magenta |
| JET | Blue | Magenta | Blue |
| PDP1 | Blue | Magenta | Magenta |
| PER | Blue | Magenta | Blue |
| SGG | Blue | Magenta | Blue |
| TIM | Blue | Magenta | Magenta |
| VRI | Blue | Magenta | Magenta |

The LRT comparing models M7 and M8 revealed statistically significant evidence of positive selection in four of the eleven genes examined: *cwo, jet, per*, and *sgg* (p < 0.05). This finding suggests that these genes have experienced adaptive evolution at specific codon sites. No significant evidence of positive selection was detected in *Clk, cry, cyc, dbt, pdp1, tim*, or *vri*, which is consistent with the strong purifying selection observed in the sliding window analysis. The genes with significantly higher LRT results are *cwo, jet, per*, and *sgg*. This supports the sliding windows result, further indicating these regions have undergone positive selection. The PAML analysis also revealed significant site-specific positive selection in three circadian clock genes: *cwo* (G203), *per* (G909), and *sgg* (G442; S452). The latter site is particularly important since it serve as an additional serine phosphorylation site of SGG in *D. melanogaster*, which are known to be important for circadian function [30]

### 3.4 Pleiotropy and Protein Evolution

Since some of the clock proteins are known to be pleiotropic, we have sought to explore the link between pleiotropy and protein divergence. To that end, we have analysed 429 additional, randomly selected proteins from the same 65 Drosophila species that were used for the clock protein analysis. The annotation and alignment of these proteins were carried out in the same way that was done with the clock proteins. Using the number of GO terms associated as a measure of pleiotropy underscores this property among clock proteins (Figure 4). Six clock proteins were amongst the top 10% of the number of associated GO terms, with PER and SGG being the most pleiotropic. Although these two proteins also exhibit elevated non-synonymous substitution rate (Ka), we did not find a significant link between Ka^*^ and pleiotropy in clock proteins, or generally among the large sample of proteins (p = 0.21, Figure 4). The gene-wise Ka also did not show a significant association with the pleiotropic score (Supplementary Figure S2). We have also estimated pleiotropy by the number of interactions that a gene/protein is known to have. Here, the linear model was marginally significant (F_1,388_=3.55, p = 0.06), but the adjusted R^2^ was low (0.006), suggesting again that protein interactions as a pleiotropy measure does not predict the level of Ka^*^ (Supplementary Figure S2).

**Fig. 4.**
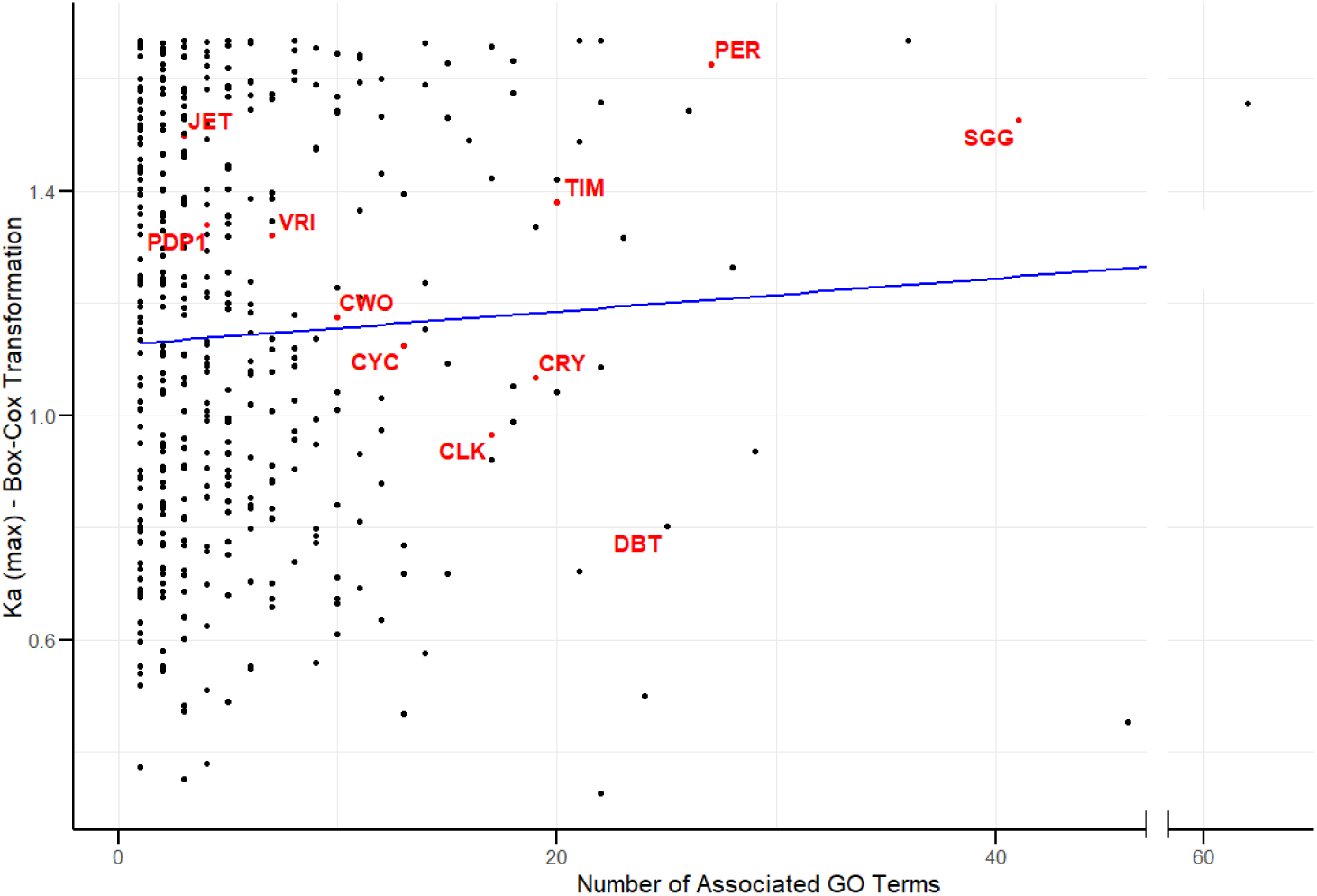
Relationship between evolutionary rate and pleiotropy in *Drosophila* proteins. The nonsynonymous substitution rate (Ka*) was calculated across 440 proteins from 65 *Drosophila* species and plotted against each protein’s pleiotropy score, measured as the number of associated GO terms. Each point represents an individual protein, with circadian clock proteins (n=11) highlighted in red. The blue line shows a linear regression fit to the data, which revealed no significant correlation between Ka and pleiotropy score (see text). Note that the x-axis contains a break to optimize the display of the data distribution.

### 3.5 Incongruency of Protein Phylogenies

Incongruency between phylogenetic trees of proteins that are members of the same networks can provide valuable insights into the evolution and co-evolution of these proteins. Here, we generated mirror-trees (tanglegrams) of different pairs of clock proteins and assessed the level of incongruency between the dendrograms. The PER-TIM tanglegram shows a substantial incongruency (Figure 5). For example, In the TIM phylogeny, the *virilis* species group (e.g. *D. virilis, D. americana, D. littoralis*) clusters with the *repleta* group (e.g. *D. repleta, D. mojavensis*), as expected for the subgenus *Drosophila*. However, the PER tree breaks this clade up. The *virilis* lineage is placed apart from the *repleta* group in the PER tree (closer to the Sophophora lineage), disrupting the monophyly of the subgenus *Drosophila*. In other words, PER’s phylogeny fails to unite the *virilis* and *repleta* clades (a deviation previously noted in smaller datasets [31]).

**Fig. 5.**
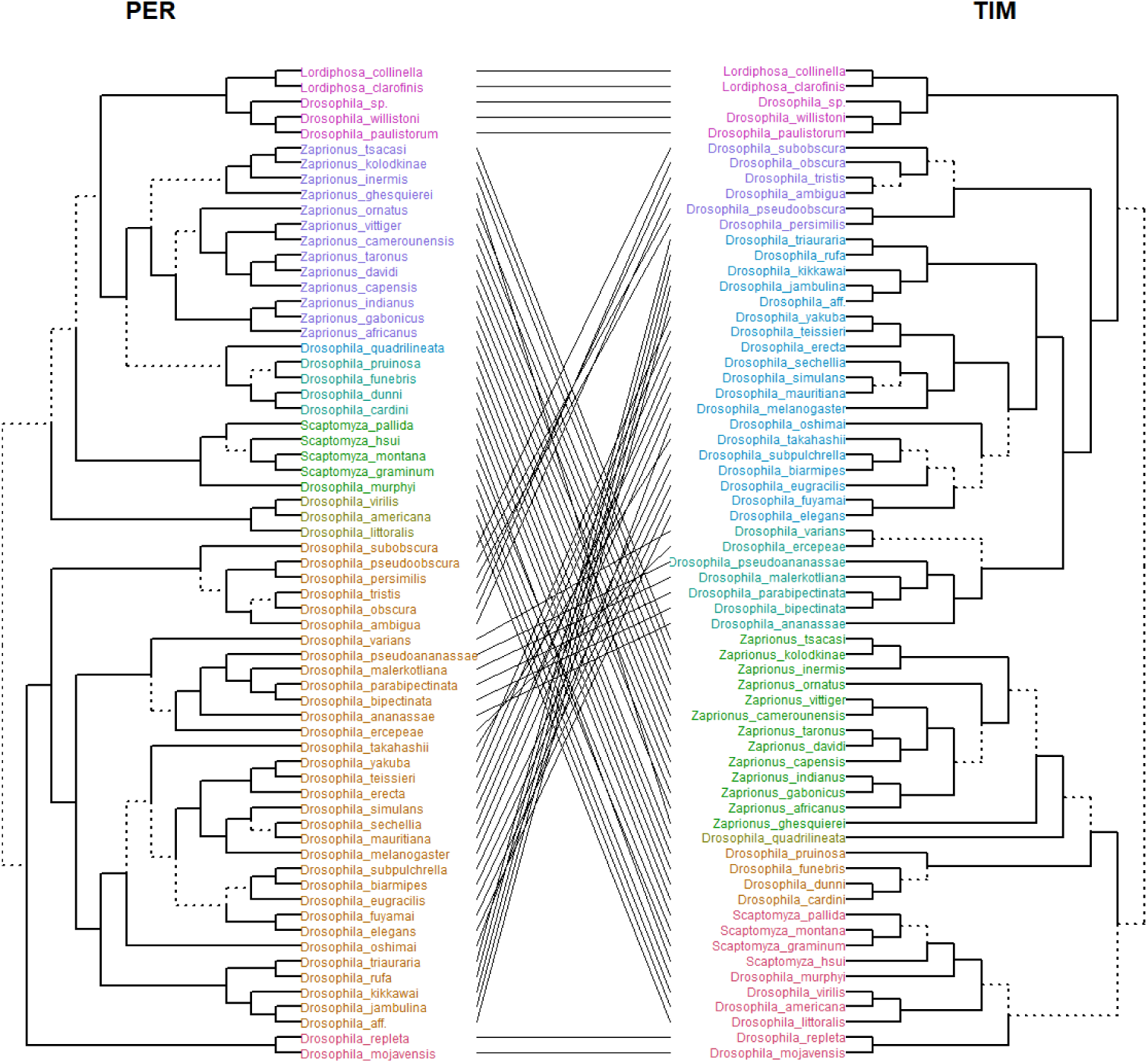
Phylogenetic incongruence between PER and TIM protein trees in *Drosophila*. To examine potential co-evolution between these core circadian clock components, we compared phylogenetic trees inferred from nucleotide alignments of PER and TIM across 65 *Drosophila* species. The mirrored trees (tanglegram) reveal multiple topological incongruences—indicated by crossing lines—suggesting differing evolutionary pressures or rates of change acting on PER versus TIM. Branches leading to distinct subtrees are marked by dashed lines.

The PER-CRY tanglegram also revealed several major topological incongruences (Supplementary Figure S3). In the CRY tree, species from the *virilis* and *repleta* groups form a single clade, consistent with their classification within subgenus *Drosophila*. In contrast, these groups are split in the PER tree, indicating disrupted monophyly. Another major difference involves the *elegans–eugracilis* lineage, which forms a clade in the PER tree but is broken in the CRY tree, where *D. elegans* is grouped with the *suzukii* subgroup instead. The relative positioning of major Sophophora subgroups (e.g., *melanogaster, ananassae, montium*) also differs, with the CRY tree generally preserving subgroup integrity while the PER tree clusters subgroups differently. These inconsistencies, especially where clades are clearly broken in one tree but intact in the other, suggest that PER and CRY have followed distinct evolutionary paths in some lineages, possibly reflecting different functional constraints or adaptive pressures. Substantial incongruencies are also present in the CLK-CYC, CLK-CWO and CRY-TIM tanglegrams (Supplementary Figure S4-S6).

## Discussion

In the current study we sought to measure the evolutionary rates that shapes the circadian clock network. Focusing on coding DNA sequences, our implicit hypothesis was that species-specific adaptations to local environmental conditions will be mirrored by the pattern of molecular divergence and substitution rate that various core clock genes undergo.

Previous studies showed that variation in a single clock gene may modify circadian function. For example, diel locomotor and sexual activity profiles differ between *D. melanogaster* and *D. psuedoobscura*, and these differences are solely due to a species variation in the *per* gene as was demonstrated by comparing the behaviour of transgenic *D. melanogaster* flies that harbored either the hetero- or the conspecific *per* gene [13]. However, in a different study, the rescue of the circadian function of *per* mutant *D. melanogaster* by different species *per* transgene yielded different levels of rescue, and this was explained by PER-TIM co-evolution [32]. Later experiments with two heterospecific transgenes expressing PER and TIM together in the double mutant *D. melanogaster* flies provided a weak or partial evidence for co-evolution [31]. Similarly, TIM transgene from remotely related species *D. ananassae* was able to rescue the circadian rhythm of D. *melanogaster* mutants [33], alluding to a weak constraint of co-evolution of TIM with other network proteins. These experiments are consistent with our finding here of modest molecular co-evolution between network proteins, which was notably not evident in pairs that are physically interacting such as PER and TIM, or CLK and CYC (Figure 2).

We acknowledge that the genetic variation in the core clock genes may capture only a small amount of the variation that drives local adaptation. Variation in other clock-controlled genes may contribute to variation in the circadian phenotypes such as the phase as was demonstrated in studies in *D. melanogaster* [34,35], or in GWAS in human populations [36]. Furthermore, some molecular adaptations in non-clock genes may modify the neuronal wiring of the clock neurons in the brain during development (rather than the cell-autonomous TTFL circuit). Species-specific wiring variations may lead to different neurochemistry of the circadian network, which may underlie the species local adaption. For example, a study comparing *D. ezoana* and *D. littoralis*, from Northern Europe, to *D. melanogaster*, an equatorial strain, revealed expression patterns of PDF and CRY in clock and non-clock neurons that correlated with the different circadian behaviour of high- and low latitudinal species [37].

Circadian clock genes are notable for their pleiotropic roles, as many of them function not only within the core timekeeping mechanism but also participate in diverse biological processes such as development, metabolism, and behavior. It is well established that genes with pleiotropic roles (influencing multiple traits or processes) are often subject to stronger purifying selection, resulting in slower sequence evolution [38,39]. The rationale is that mutations in a highly pleiotropic gene can have deleterious effects on numerous traits, imposing a “cost of pleiotropy” that constrains divergence [39]. Prior studies across diverse systems support this expectation. For example, genomewide analyses in yeast showed that genes affecting many phenotypes tend to be more evolutionarily conserved than those with narrow effects [38]. Similarly, in complex networks, core hub genes (which are often pleiotropic) exhibit reduced protein evolutionary rates compared to peripheral genes. In a co-expression network of European aspen, highly connected core genes showed significantly lower sequence divergence, reflecting strong selective constraint [40]. Similarly in vertebrates, comparative studies have noted that pleiotropic genes can slow the pace of evolutionary change across species, underscoring their essential, constrained nature in development and physiology (e.g., [41]). These prior findings foster a general expectation that pleiotropic genes – such as many circadian clock components – should evolve under stringent constraint and accumulate fewer amino acid substitutions over time.

In light of these expectations, we specifically examined whether the circadian clock genes’ known pleiotropy correlates with heightened evolutionary constraint. Many core clock proteins in *Drosophila* have well-documented roles beyond timekeeping (e.g., involvement in metabolism, development, or other signaling pathways), making them classic candidates for pleiotropic constraint. Surprisingly, our analysis did not support the anticipated pleiotropy–divergence relationship. Despite using multiple metrics to quantify gene pleiotropy, we found no significant correlation between a gene’s pleiotropic degree and its rate of protein sequence evolution (non-synonymous substitution rate, *K*a) across 65 *Drosophila* species. Highly pleiotropic clock genes such as *sgg* or *Pdp1*, which might be expected to evolve especially slowly, did not show consistently lower divergence than less pleiotropic clock genes. In fact, the circadian genes with extensive extra-circadian functions were not universally more conserved at the sequence level than their more specialized counterparts. This result stands in contrast to the strong evolutionary constraints predicted by the “cost of pleiotropy” model and reported in other biological networks [39]. Our findings align with recent observations that the impact of pleiotropy on evolutionary rate can be more nuanced than initially assumed; in some cases, highly pleiotropic genes do not exhibit markedly slower evolution, indicating that pleiotropy alone is not an absolute determinant of sequence conservation [42].

The lack of a pleiotropy–divergence correlation in the circadian system is notable given prior evidence from plants, animals, and fungi that pleiotropy constrains molecular evolution. For instance, a study in *Populus* found that genes with high network connectivity (a proxy for pleiotropy) had significantly lower nonsynonymous divergence, consistent with stronger purifying selection [40]. Likewise, He and Zhang’s [38] yeast study demonstrated a positive correlation between the number of traits a gene influences and its evolutionary conservation.

Our analysis of the circadian clock genes did not recapitulate these patterns. Even though several clock genes act as network “hubs” interfacing with multiple pathways (and might thus be considered indispensable), they have not been uniformly constrained in their coding evolution. This discrepancy suggests that the relationship between pleiotropy and evolutionary rate may depend on biological context or the nature of the pleiotropic effects. It is possible that circadian proteins, despite their multi-functionality, contain modular domains or separate interaction interfaces that mitigate the fitness costs of amino acid changes. If different functions of a pleiotropic protein are compartmentalized into distinct regions of the molecule, certain regions can diverge without compromising all functions, thereby relaxing the overall constraint on sequence evolution. Such modular pleiotropy could allow clock proteins to accumulate changes in specific sites (for example, those affecting circadian period tuning or protein–protein interactions within the clock) while preserving other crucial functions via conserved domains.

Our findings highlight that the circadian clock network, despite its interwoven architecture and pleiotropic components, does not enforce an unusually slow evolutionary rate on its genes. This contrasts with a simplistic view that pleiotropy invariably slows evolution, and instead hints at an inherent flexibility in how clock genes balance multiple roles. One interpretation is that adaptive evolution in clock genes might occur through mechanisms other than protein-coding change—such as cis-regulatory modifications or gene expression adjustments—thereby sidestepping the pleiotropic constraints on the protein sequence. Zhang [43] noted that evolutionary adaptation can sometimes be channeled through regulatory changes in pleiotropic genes to avoid detrimental effects on other traits. In the circadian system, species-specific adaptations of rhythms could predominantly involve changes in gene regulation or network interactions, allowing core clock protein sequences to drift more freely than expected. Additionally, our broad analysis (covering hundreds of *Drosophila* genes) suggests that pleiotropy per se is not a straightforward predictor of evolutionary rate in this lineage. This supports the idea that while pleiotropic genes are often under strong purifying selection, their evolutionary trajectories can be modulated by factors like protein structure, network redundancy, and the mode of pleiotropy (e.g. whether a gene’s multiple functions are tightly linked or can vary independently).

In summary, the circadian clock genes provided an empirical test of the long-held assumption that pleiotropy constrains molecular evolution. Contrary to expectations from earlier studies and theoretical models, we found no evidence that pleiotropic clock genes evolve more slowly at the sequence level. This divergence from prior patterns underscores the complexity of evolutionary dynamics in protein networks. It suggests that pleiotropy’s impact on sequence divergence is context-dependent and may be mitigated by modular functionality or alternative adaptive pathways. Our results contribute to a growing recognition that gene network evolution can accommodate pleiotropic versatility without universally imposing strong evolutionary constraint, thereby allowing even multifunctional proteins like clock components to explore sequence space and potentially facilitate lineage-specific innovations in circadian biology.

