## Supplementary Figure for "The Evolutionary Flexibility of the Drosophila Circadian Clock: Network Constraints or Adaptive Freedom?"

**This PDF file includes:**

Supplementary Figures S1 to S6

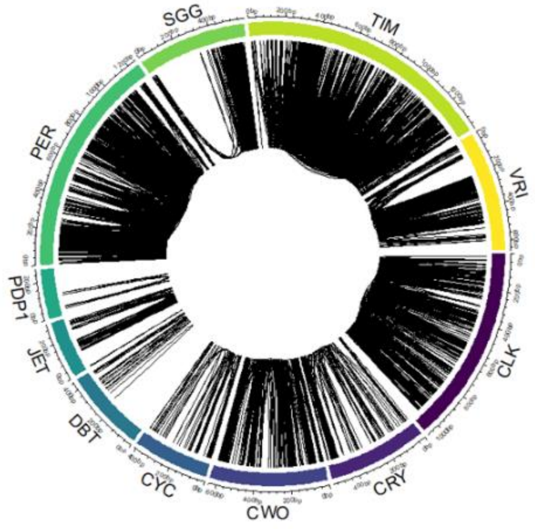


**Figure S1. Co-evolving sites within each circadian clock protein.** The circular plot shows significant co-evolving pairs of amino acids ($p < 0.05$).

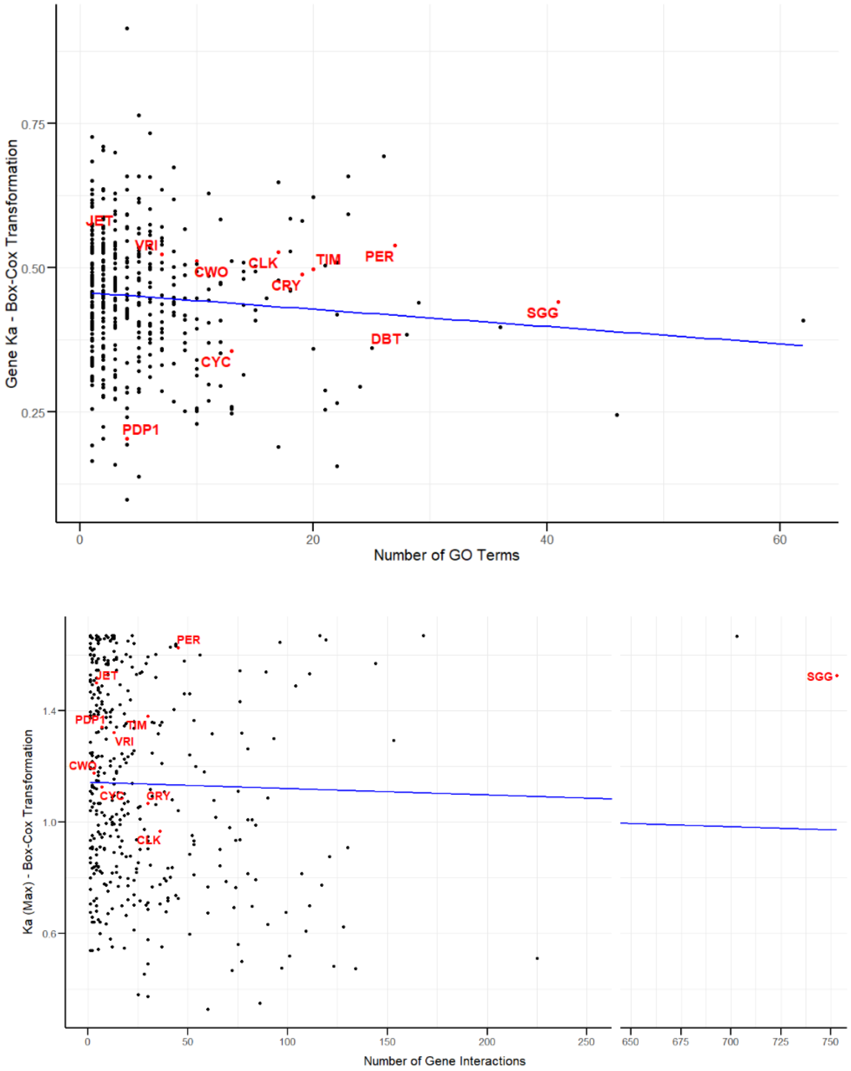


**Fig. 4. Relationship between evolutionary rate and pleiotropy in *Drosophila* proteins.** The nonsynonymous substitution rate (Ka*) was calculated across 440 proteins from 65 *Drosophila* species and plotted against each protein's pleiotropy score, measured as the number of associated GO terms (Top) and the number of interactions that a gene/protein is known to have (Bottom). Each point represents an individual protein, with circadian clock proteins (n=11) highlighted in red. The blue line shows a linear regression fit to the data, which revealed no significant correlation between Ka and pleiotropy score (see main text). Note that the x-axis in the bottom graph contains a break to optimize the display of the data distribution.

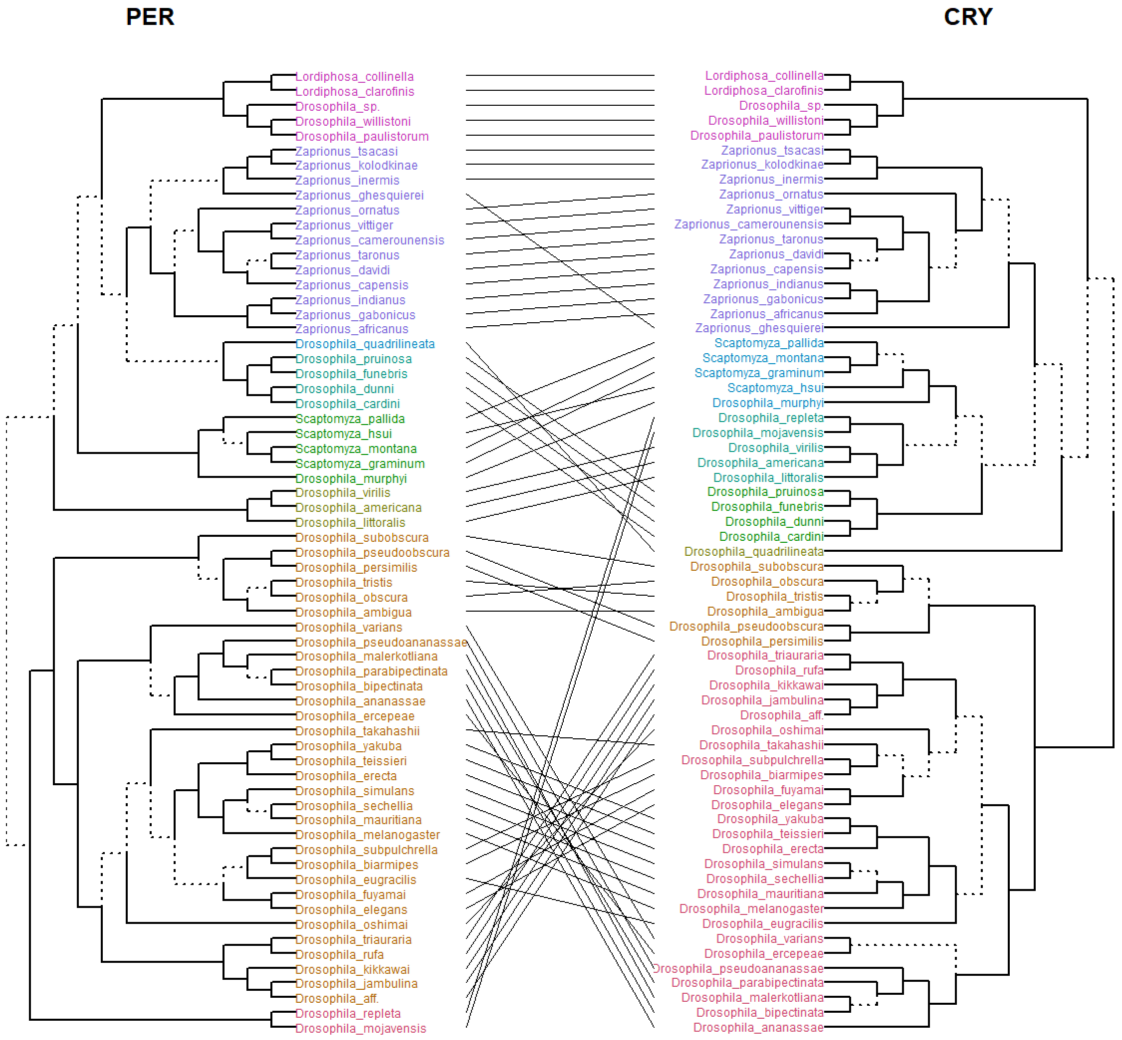


Figure S3. **Phylogenetic incongruence between PER and CRY protein trees in *Drosophila***. To examine potential co-evolution between these core circadian clock components, we compared phylogenetic trees inferred from nucleotide alignments of PER and CRY across 65 *Drosophila* species. The mirrored trees (tanglegram) reveal multiple topological incongruences—indicated by crossing lines—suggesting differing evolutionary pressures or rates of change acting on PER versus CRY. Branches leading to distinct subtrees are marked by dashed lines.

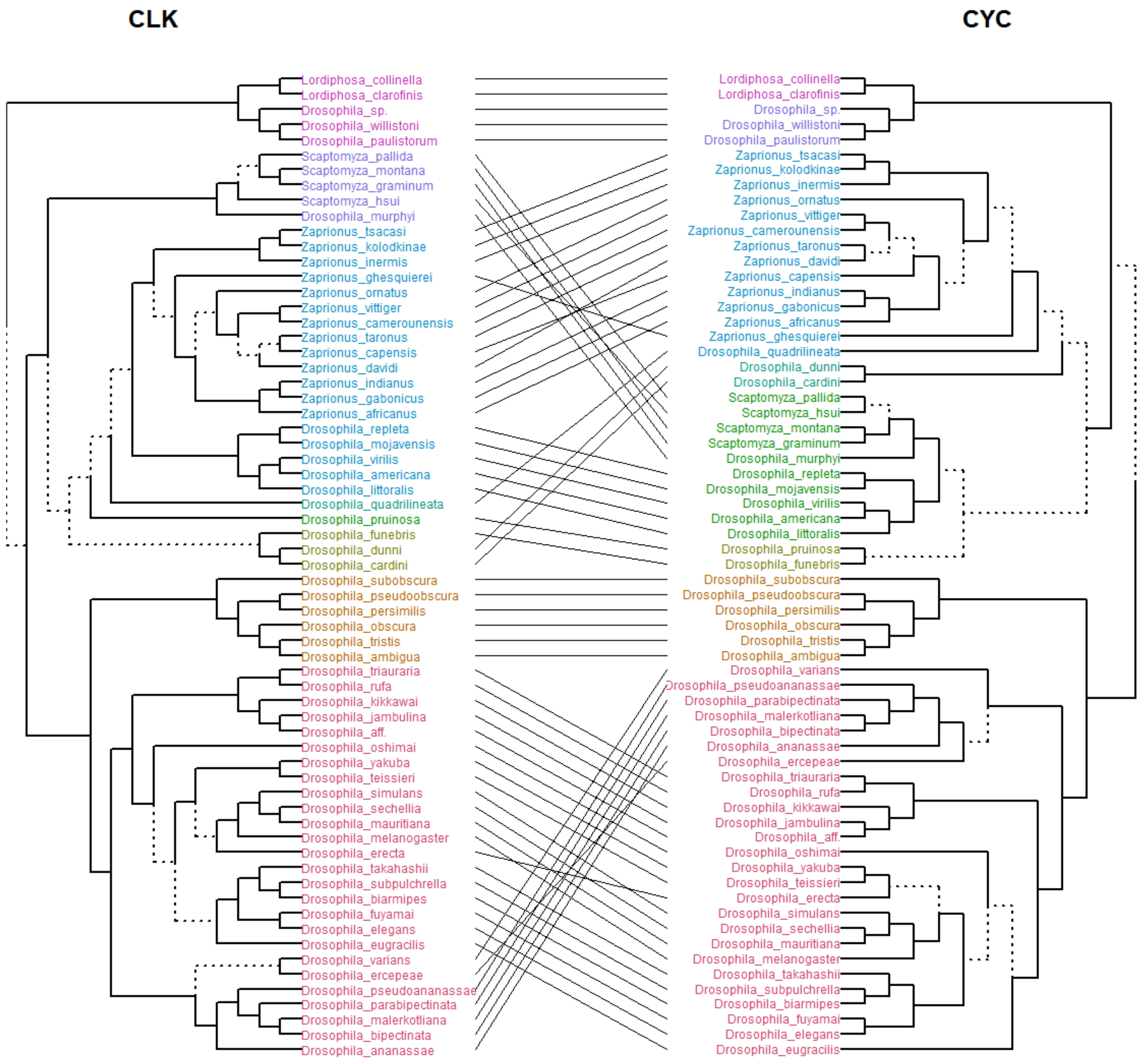


Figure S4. **Phylogenetic incongruence between CLK and CYC protein trees in *Drosophila*.**

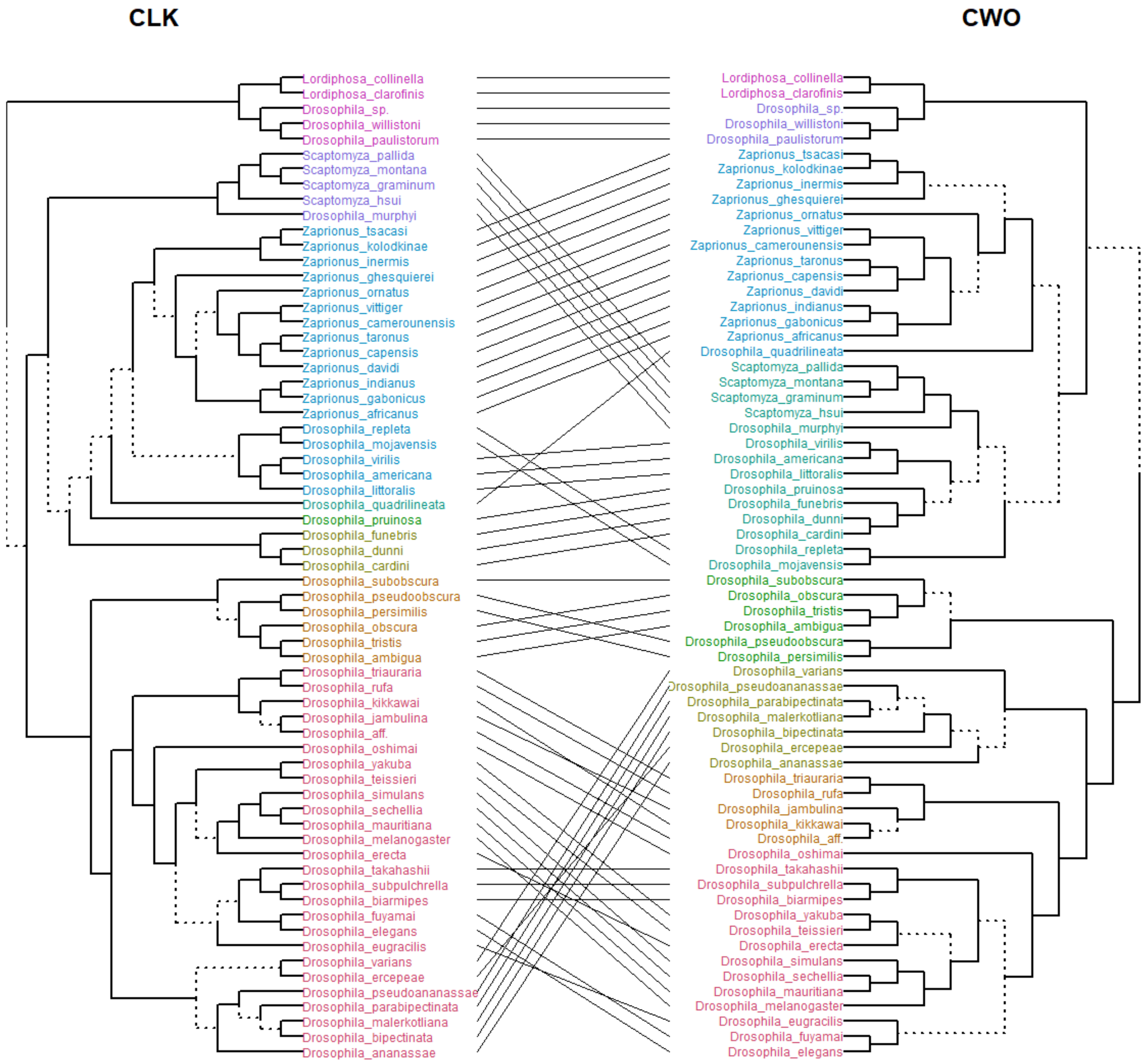


Figure S4. **Phylogenetic incongruence between CLK and CWO protein trees in *Drosophila*.**

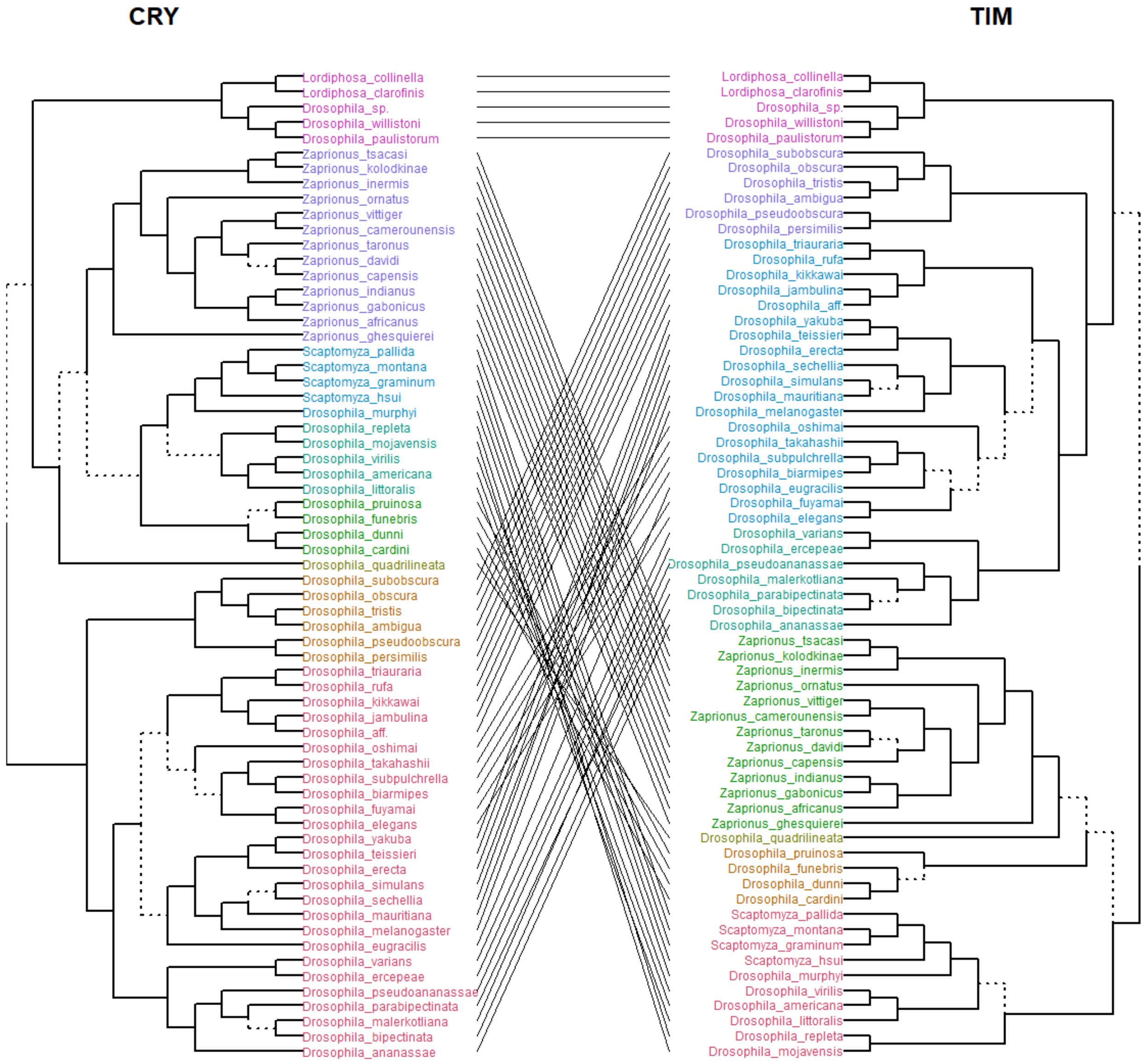


Figure S5. **Phylogenetic incongruence between CRY and TIM protein trees in *Drosophila*.**

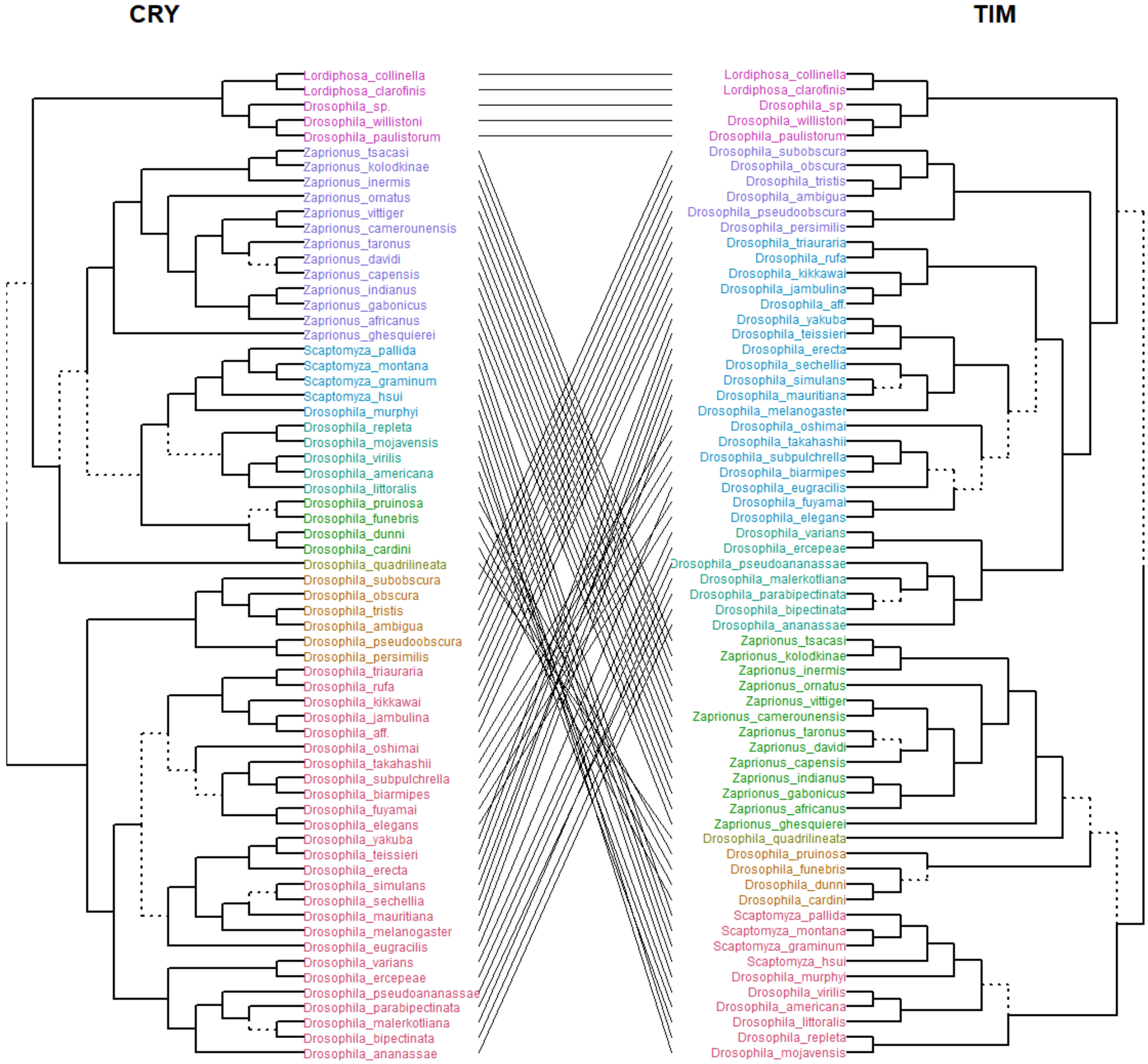


Figure S6. **Phylogenetic incongruence between CRY and TIM protein trees in *Drosophila*.**
